# Arteria: An automation system for a sequencing core facility

**DOI:** 10.1101/214858

**Authors:** Johan Dahlberg, Johan Hermansson, Steinar Sturlaugsson, Pontus Larsson

## Abstract

Arteria is an automation system aimed at sequencing core facilities. It is built on existing open source technologies, with a modular design allowing for a community-driven effort to create plug-and-play micro-services. Herein we describe the Arteria system and elaborate on the underlying conceptual framework. The Arteria system breaks down into three conceptual levels; orchestration, process and execution. At the orchestration level it utilizes an event-based model of automation. It models processes, e.g. the steps involved in processing sequencing data, as workflows and executes these in a micro-service based environment. This creates a system which is both flexible and scalable. The Arteria Project code is available as open source software at http://www.github.com/arteria-project.

## Introduction and background

Nucleotide sequencing is the practice of determining the order of bases of the nucleic acid sequences that form the foundation of all known forms of life. It has been hugely successful as a research tool, used to understand basic biology [1–3], and is also applied as a tool for precision medicine [4]. Major technological advances during the last decade have enabled high throughput approaches for massively parallel sequencing (MPS) [5]. The amount of data generated globally from MPS has boomed in recent years, and has been projected to reach a yearly production of 10^21^ base-pairs per year by 2025, demanding 2-40 Exa-bytes (10^18^) per year of storage [6]. This massive expansion places new demands on how data is analyzed, stored and distributed.

A large amount of this nucleotide sequencing is carried out at sequencing core facilities, which perform sequencing as a service. Exactly which services are provided vary widely, but typically delivery of raw sequencing data after conversion to a standard fastq format is a minimum [7].

Automation of both the laboratory and computational procedures is crucial in order for a sequencing facility to be able to scale with respect to the amount of samples processed. Furthermore, automated processes reduce the risk of human errors, which contributes to higher quality data. A challenge in this context is that despite the fairly standard lab protocols, small changes in procedures, infrastructure and surrounding systems create a combinatorial situation that makes every lab unique. Most sequencing core facilities have developed their own bespoke solutions to this problem, and these are often highly coupled to the exact infrastructure and process of that particular core facility [8].

In recent years there has been an increased interest in workflow systems, both in academia [8–11] and in industry [12,13]. Typically these systems address the issue of modelling a workflow as a directed acyclical graph of dependencies between computational tasks, which can then be executed. These are often designed to be run on a per project or per sample level, with parameters being provided manually by the operator. This model is well suited for processing large amounts of data, where all samples in a project can be analyzed more or less the same way. However for institutions that provide sequencing as a service to many users or projects, this type of system does not typically scale well, due to the need for manual intervention at different stages of the process.

Furthermore, there are additional aspects associated with operating a sequencing facility that are not addressed by these types of systems. Examples of this include automatically starting processing of data as a sequencing run finished, archiving of data to remote storage and removal of data once certain criterias are met. These *operational* aspects have not been as thoroughly investigated in the scientific literature, but are essential when taking a bird’s-eye view of the complete process of refining raw MPS data to scientific results on a high-throughput scale.

Tackling these issues also involves examination of how higher level orchestration integration and management of such workflows can be done in an efficient yet flexible manner, while providing a clear enough understanding of the system so that changes can be implemented with minimal mental overhead and risk of breaking existing functionality. One example of a system addressing this problem in the context of a sequencing core facility is described by Cuccuru et. al [14]. They describe a system with a central automator that handles orchestration of the processes in an event-based manner, utilizing a separate workflow manager.

Herein, we describe the automation system Arteria, which is available as open source software at: https://github.com/arteria-project. Arteria utilizes the open source automation platform StackStorm [15] for event-based orchestration, the Mistral [16] workflow engine for process modelling, and micro-services for action execution. Arteria has been successfully implemented for sequencing data processing at the SNP&SEQ Technology Platform, where it has been instrumental in scaling up operations to meet a large increase in sequencing capacity, going from 187 Tbases in 2015 to 490 Tbases in 2016, the year in which Arteria was brought into production usage.

## Definitions

| Term | Description |
| --- | --- |
| Action | A computational unit of work, e.g. processing a file or inserting data into a database. This is sometimes referred to as a task. |
| Process | A set of steps that have to be finished to achieve a particular goal, e.g. delivering data to a user. A process can include automated and manual steps. |
| Workflow | A workflow models a process, as a number of *actions* following each-other. This can be described by a directed acyclic graph. |

## System overview

The Arteria system is built with two existing open source technologies at its core; the StackStorm automation platform [15] and the Mistral workflow service [16]. By adopting existing open source solutions and extending them for our domain we are able to leverage the power of a larger open source community. This has allowed us to focus on our specific use-case; automation of our sequencing data processing.

The Arteria system can be divided into three conceptual levels, a model that has been adopted from StackStorm: the orchestration level, the process level and the execution level (figure 1). At the highest level, the orchestration level, StackStorm serves as the central point of automation. It utilizes an event-based model to decide when actions should be triggered. An example of an event that should trigger actions to be taken by the system can be a sequencing run finishing. In addition,it provides command line and web-based user interfaces through which an operator can interact with the system.

**Figure 1.**
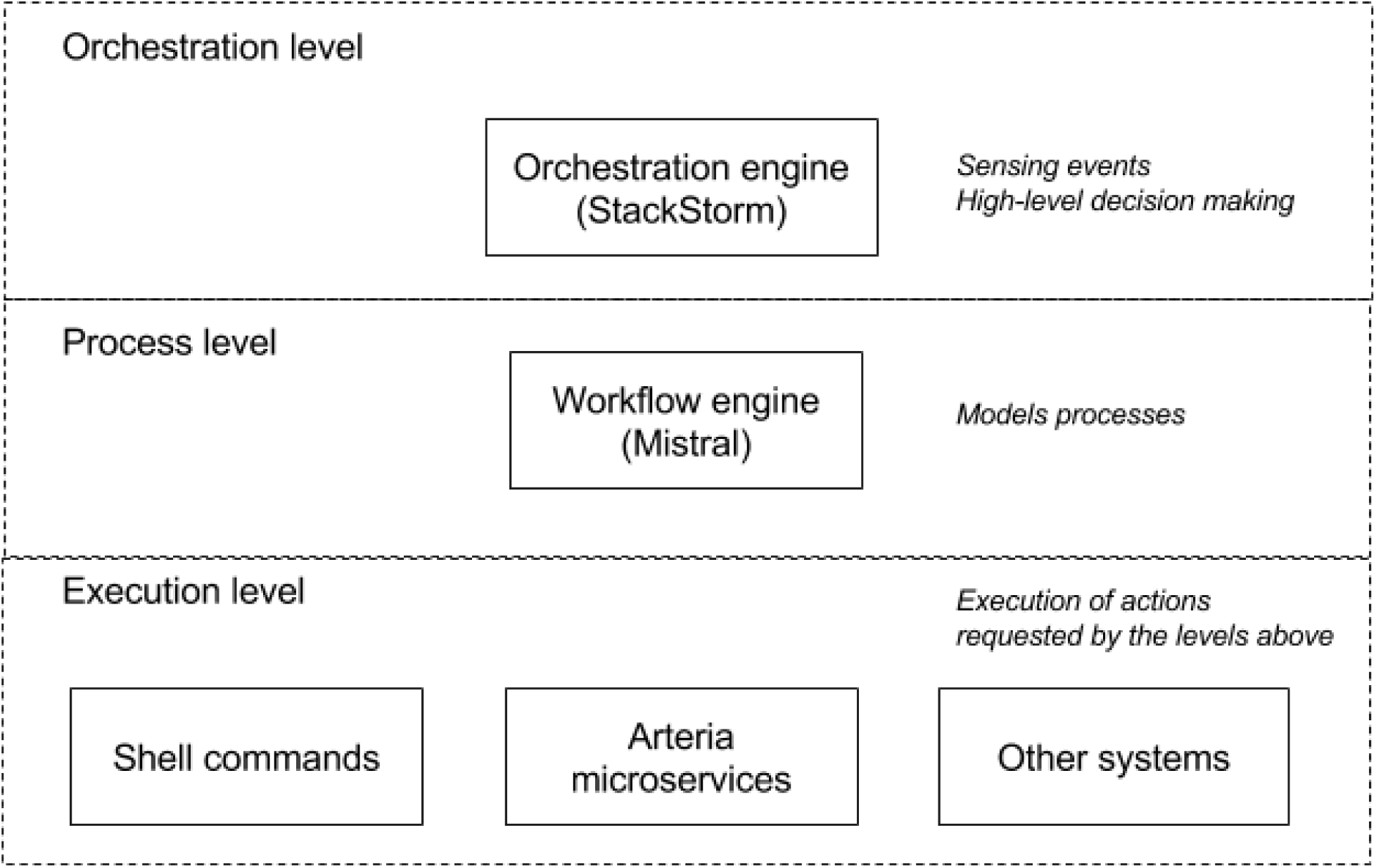
An overview of the conceptual levels of the Arteria project.

At the process level our internal processes are modelled as workflows using the Mistral workflow service. An example of such a workflow is the one which takes raw data once the sequencing instrument is finished, carries out basic processing, gathers quality control data and transfers the data to a high-performance computing resource.

Finally, at the execution level, actions are carried out. This level includes multiple modes of execution, ranging from running a shell command on a local or remote machine, to interacting with surrounding systems such as a laboratory information management system (LIMS) or issuing a command to a micro-service. The final mode, the micro-service, is the one favoured by Arteria. It has the advantages of making the system flexible and decoupling details of an execution from the process in which they take part.

This separation of the system into levels makes the Arteria system easier to reason about, and places implementational details at the correct level of abstraction. In addition to this, Arteria enforces a separation of concerns that makes it easier to update or replace individual components, without having to make large changes to the system as a whole. This creates a flexible system which is able to meet the demands on scaling placed on sequencing core facilities, where protocols are modified and new instrumentation is implemented constantly to meet the the users needs.

### Event-based orchestration

At the orchestration level we use StackStorm to coordinate tasks. A core concept of StackStorm is its event-based model of automation (see figure 2). It utilizes sensors to pick up events from the environment. A typical example of this is a sequencing instrument finishing a run. The event is then passed through a rule layer that decides which, if any, action should be taken given the parameters of the event. This simple yet powerful abstraction makes the Arteria system and its behaviour simple to reason about. In addition to triggering actions in response to sensor events, an operator can manually initiate an action either via a command line or web interface.

**Figure 2.**
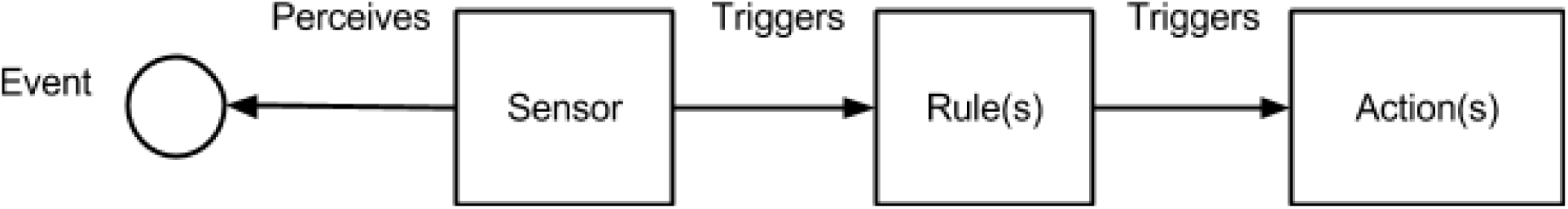
Description of the StackStorm event model. Sensors will perceive events in the environments, e.g. a file being created or a certain time of day it occurs. This passes information to the rule layer where the data is evaluated and depending on which, if any, criteria are fulfilled one or more actions are triggered. Actions can be single commands or full workflows to be executed.

Furthermore StackStorm provides per-action monitoring capabilities. Each action taken by the Arteria system is assigned a unique id, allowing operators to follow the progress of processes in the system. An additional advantage of this is that it can be used to create the audit-trails, which are one of the components required in the European quality standard ISO/IEC 17025 [17] under which the SNP&SEQ Technology Platform operates. Finally, providing a centralized interface to the underlying processes means that the number of systems to which operators need explicit shell access is reduced, which is an advantage from a security perspective. Additionally, Arteria forms an abstraction level to the underlying systems lowering the knowledge requirements in e.g. linux systems for the operator.

### Modelling processes as workflows

At the process level the process of a particular use case is modeled using the Mistral workflow language. For example, this can mean translating documentation of an existing process to a workflow, thus reducing the amount of manual work required, as well as reducing the risk of human errors. Mistral uses a declarative yaml syntax to define a workflow, which allows for the definition of relatively complicated dependency structures. It supports the use of conditionals, forking and joining. It will execute actions that do not have dependencies on each other concurrently. This simple and powerful syntax has the advantage of also serving as a human-readable documentation of the modelled process.

### Micro-services provide a flexible execution model

Finally we have the execution level. At this level any action that needs to be carried out by the system is actually executed. In this case Arteria favors the use of single-purpose microservice executors, and these provide the actual functionality and logic for performing the actions. These micro-services are called from the process level via an HTTP API, making the communication simple and allowing for easy integration with other services. An example of such a microservice is the one provided by Arteria to run the preprocessing program Illumina bcl2fastq [18], which processes the raw data produced by an Illumina sequencing instrument and converts it to the industry standard fastq-format. However, this approach is flexible enough that it can also include running a shell command or calling and updating another service, e.g. a LIMS.

Using micro-service as the primary execution mode increases the flexibility of the Arteria system as the implementational details of *how* something is run is decoupled from *when* it is run. Furthermore it means that such micro-services can be reused across systems, or even centers, creating an avenue for reuse and collaboration, which sets the Arteria approach apart from other sequencing core facility systems that are typically tightly coupled to the process and infrastructure of the sequencing core facility that developed it.

Finally, decoupling the execution layer has allowed us to build simple interfaces for existing softwares included under the ISO/IEC 17025 standard accreditation, thus significantly reducing the burden of having to reimplement softwares that have been used reliably for a long time in operation.

## Deployment scenario and usage statistics

At the SNP&SEQ Technology Platform sequencing core facility, the Arteria system is deployed in a distributed environment (see figure 3) and orchestrates actions across a local cluster of 10 nodes used for storage and preliminary analysis with 208 cores and 120 TB of storage capacity, as well as a high-performance computing cluster at the Uppsala Multidisciplinary Center for Advanced Computational Science (UPPMAX) high-performance computing center with 4000 cores and 1.1 PB storage. This system is fully capable of supporting the fleet of 10 Illumina sequencers (5 HiSeqX, 2 HiSeq2500, 1 MiSeq, and 1 NovaSeq) which are currently in use at the SNP&SEQ Technology Platform.

**Figure 3.**
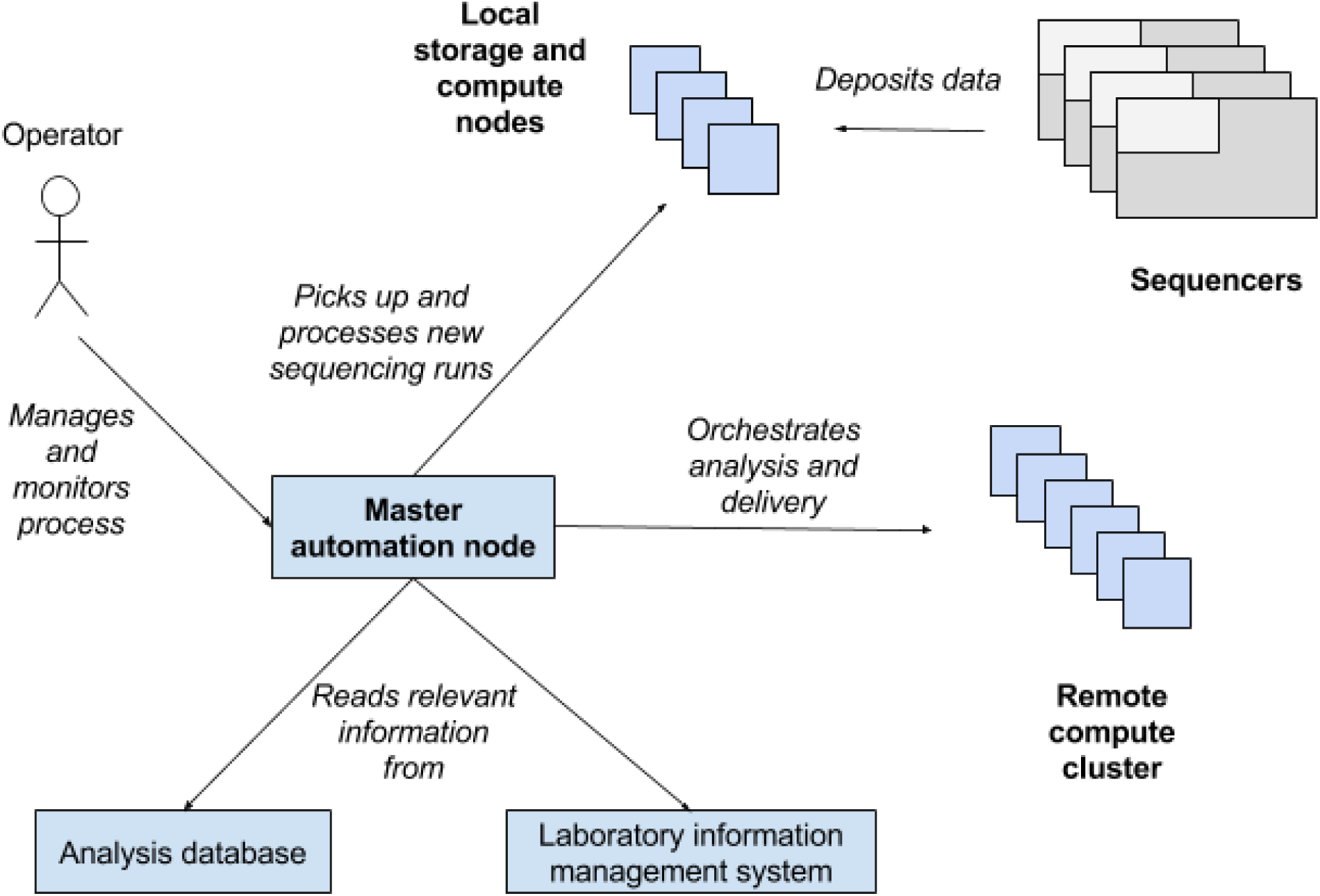
Schematic view of a system deployment scenario, showing how data is written to the local storage and compute nodes from the sequencing machines, and how the system uses information and resources from multiple sources to coordinate the process. The operator can then monitor and control the processes from the single interface provided at the master automation node.

Since being deployed at the SNP&SEQ Technology Platform, the Arteria system has been used to process more than 22000 samples and 326 projects, which corresponds to ~640 Tera-bases of sequencing data. We have been able to update our process at regular intervals, which is shown by the fact that there has been 30 releases of the code describing our workflows, since it was deployed into production.

## Discussion

In this paper, we describe the automation system Arteria, which is built on top of the StackStorm automation platform and the Mistral workflow service. While the Arteria system has been in production at the SNP&SEQ Technology Platform sequencing core facility we have increased our capacity by a factor two and the system has been a significant factor in allowing us to increase our throughput in terms of projects, samples and data.

Arteria presents an approach to managing the full breadth of the operational aspects surrounding sequencing center operations. It manages *when* as well as *how* certain processes are to be carried out. Through the use of StackStorm as the orchestration engine, we are able to both have a framework for the development of new functionality as well as providing a unified user interface to the system operators.The use of workflows at the process level, through Mistral, reduces the need for additional documentation and lowers the risk of human errors. Furthermore, the use of workflows allows for changes to the process to be code reviewed, in accordance with best practices in software development. Finally, the use of micro-service at the execution level has enabled a greater degree of flexibility in the execution model, a clear separation of responsibilities between services, as well as the integration of existing software. Being able to easily integrate existing software into the system has enabled quicker implementation as it lowers the burden of validation for the ISO/IEC 17025 standard accreditation.

Arteria takes advantage of existing open source tools and aims at creating an avenue for collaboration between sequencing core facilities. We believe that decoupling process from execution, especially the micro-services developed within the Arteira project, could serve as fertile ground for collaboration. The stand-alone nature of the micro-services means that it should be possible for anyone interested to pick them up and include them into their own operations.

We recognize that this type of approach has a higher initial overhead then e.g. an orchestration system based on scripts and cron-tab entries. This overhead is payed for both in terms of additional hardware requirements (current production hardware requirements are: a quad core CPU, >16GB RAM, 40G of storage) and increased system complexity. This increase in system complexity can make debugging more difficult. However, in the long run we are confident that the additional overhead pays off, by proving a solid and extensible framework for developing new functionality in accordance with our core facility’s needs, without requiring extensive changes to the existing infrastructure.

All components of Arteria are open source and available to the wider community (https://github.com/arteria-project). Finally, it is our hope that there are valuable lessons to be drawn from the design described here for anyone who needs to implement an orchestration system, in the context of a sequencing core facility, or elsewhere.

## Acknowledgements

This work was performed at the SNP&SEQ Technology Platform in Uppsala, which is part of the National Genomics Infrastructure (NGI) funded by Science for Life Laboratory (SciLifeLab) and the Swedish Council for Research Infrastructures. The entire bioinformatics team at the SNP&SEQ Technology Platform has contributed to the Arteria project with feedback during its adoption. Especially Monika Brandt, Matilda Åslin and Sara Ekberg have provided extremely useful feedback based on their daily operations of the system, as well as contributing code to the project. Parts of the computations for the Arteria Project were performed on resources provided by SNIC through Uppsala Multidisciplinary Center for Advanced Computational Science (UPPMAX).

We would like to acknowledge Samuel Lampa and Jessica Nordlund for reading this text and providing highly valuable comments, which have been instrumental in developing this manuscript.

